# Diel Partitioning in Microbial Phosphorus Acquisition in the Sargasso Sea

**DOI:** 10.1101/2024.03.28.587236

**Authors:** Daniel Muratore, Naomi E. Gilbert, Gary R. LeCleir, Steven W. Wilhelm, Joshua S. Weitz

## Abstract

The daily cycle of photosynthetic primary production at the base of marine food webs is often limited by the availability of scarce nutrients. According to temporal niche partitioning theory, competition for scarce resources can be alleviated insofar as the intensity of nutrient uptake and assimilation activities are distributed heterogeneously across organisms over periodic input cycles. Recent analysis of community transcriptional dynamics in the nitrogen-limited subtropical North Pacific gyre revealed evidence of temporal partitioning of nitrogen uptake and assimilation between eukaryotic phytoplankton, cyanobacteria, and heterotrophic bacteria over day-night cycles. Here, we present results from a Lagrangian metatranscriptomic time series survey in the Sargasso Sea and demonstrate temporally partitioned phosphorus uptake in this phosphorus-limited environment. In the Sargasso, heterotrophic bacteria, eukaryotic phytoplankton, and cyanobacteria express genes for phosphorus assimilation during the morning, day, and dusk, respectively. These results support the generality of temporal niche partitioning as an emergent mechanism structuring uptake of limiting nutrients and facilitating coexistence of diverse microbes in open ocean ecosystems.

## Introduction

In the ocean, communities of microorganisms set ecological rhythms of primary production and respiration in response to the daily light cycle, fueling the base of the marine food web and modulating biogeochemical cycles [7, 14, 17, 21, 35]. Diel oscillations in carbon fixation among photosynthetic plankton cascade through community metabolism and gene expression, resulting in emergent ecosystem-level cycles in biomass and biomass molecular composition [2–4, 13, 28, 29]. Diel transcriptional and metabolic organization also expands beyond carbon metabolism. Field studies in nitrogen-starved ocean regions [37] have identified diel transcriptional cycles in nitrogen uptake [39] and nitrogen fixation [40]. Heterotrophic taxa from the same region have also been reported to express organic nutrient transporters on a diel cycle [28–30]. Organisms taking turns in fulfilling specific ecological niches or taking up resources has been termed ‘temporal niche partitioning’ in terrestrial ecology [19, 31]. Niche partitioning over diel cycles has been ascribed to predatory behavior in marine animals [20] and invoked in microbial community successional dynamics in salt marsh environments [8]. Recently, asynchronous diel cycles have also been identified amongst diverse taxa in the nitrogen-stresed North Pacific Subtropical Gyre (NPSG), suggesting that differential uptake and utiluization of a limiting nutrient could enable coexistence in diverse communities [28].

The Sargasso Sea is an open ocean environment where plankton are considered to be persistently phosphorus-stressed at the surface [9, 35, 37]. In the subtropical northwestern Atlantic, concentrations of dissolved phosphate can be sub-nanomolar [41], sufficient to induce phosphate stress in primary producers [9, 37]. Diverse strategies have evolved to mitigate phosphorus limitation among marine microbes. For example, sulfono- and betaine lipids have been shown to replace phospholipids as a method of reducing cell phosphorus quotas [6, 27, 38]. Utilizing phosphonates, organic phosphorus-containing molecules, as an alternative phosphorus source has also been described in heterotrophic [34] and photosynthetic bacteria [33]. In the Sargasso Sea, dissolved organic phosphorus has been measured to average almost 95% of total dissolved phosphorus [24]. High copy numbers of the C-P lyase *phnJ* gene and strong responses in a proxy for C-P lyase activity have also been observed *in situ* [32]. This evidence reinforces the ecological relevance of phosphorus limitation in the Sargasso Sea, similar to the relevance of nitrogen limitation in the North Pacific Subtropical Gyre [28]. Indeed, field studies of the diazotrophic cyanobacterium *Trichodesmium* have found the presence of organic and inorganic phosphorus acquisition genes [11] that are expressed on a diel cycle [10] consistent with pervasive phosphorus stress in this region.

Here, we tested the hypothesis that diverse microbes in a phosphorus-limited ecosystem will partition phosphorus uptake over the diel cycle within a Lagrangian metatranscriptomic time series survey, sampling every 4 hours over the course of 5 days. Our metatranscriptomic analyses demonstrated widespread temporal partitioning of diverse phosphorus uptake and incorporation genes, suggesting a diel cascade of phosphorus uptake from heterotrophic bacteria at sunrise, to eukaryotic phytoplankton during the day, to cyanobacteria at dusk. The results reinforce the generality of temporal niche partitioning as a mechanism of competition alleviation in open ocean microbial ecosystems.

## Results

We analyzed 58 metatranscriptomes collected during a Lagrangian sampling expedition on the R/V *Atlantic Explorer* (AE1926) from October 12-17, 2019 (Figure 1a) in the vicinity of the Bermuda Atlantic Time Series (BATS) study location in the Sargasso Sea to examine diel changes in microbial community-level gene expression (SI Data S1). Using a nonparametric method with multiple testing correction set at a 10% false discovery rate [36], we evaluated whether gene transcripts had 24-hour periodicity in their abundance patterns (see Methods). In all, we evaluated 97,829 different genes from eukaryotic phyla, cyanobacterial genera *Prochlorococcus* and *Synechococcus*, and heterotrophic bacterial orders for diel periodicity. Our analysis found 11,194 significantly diel genes (SI Data S2). Diel genes spanned 21 phyla of eukaryotic phytoplankton (9.33% of total genes), the cyanobacterial groups *Prochlorococcus* and *Synechococcus* (37.4% of total genes), and heterotrophic bacteria representing 6 major classes comprised of 44 orders (13.2% of total genes). A similar analysis conducted on data from a cruise in the North Pacific Subtropical Gyre examined 64,011 genes [28], and found a similar proportion (8.94% vs 11.4% reported here) of genes with diel transcript abundances.

**Figure 1.**
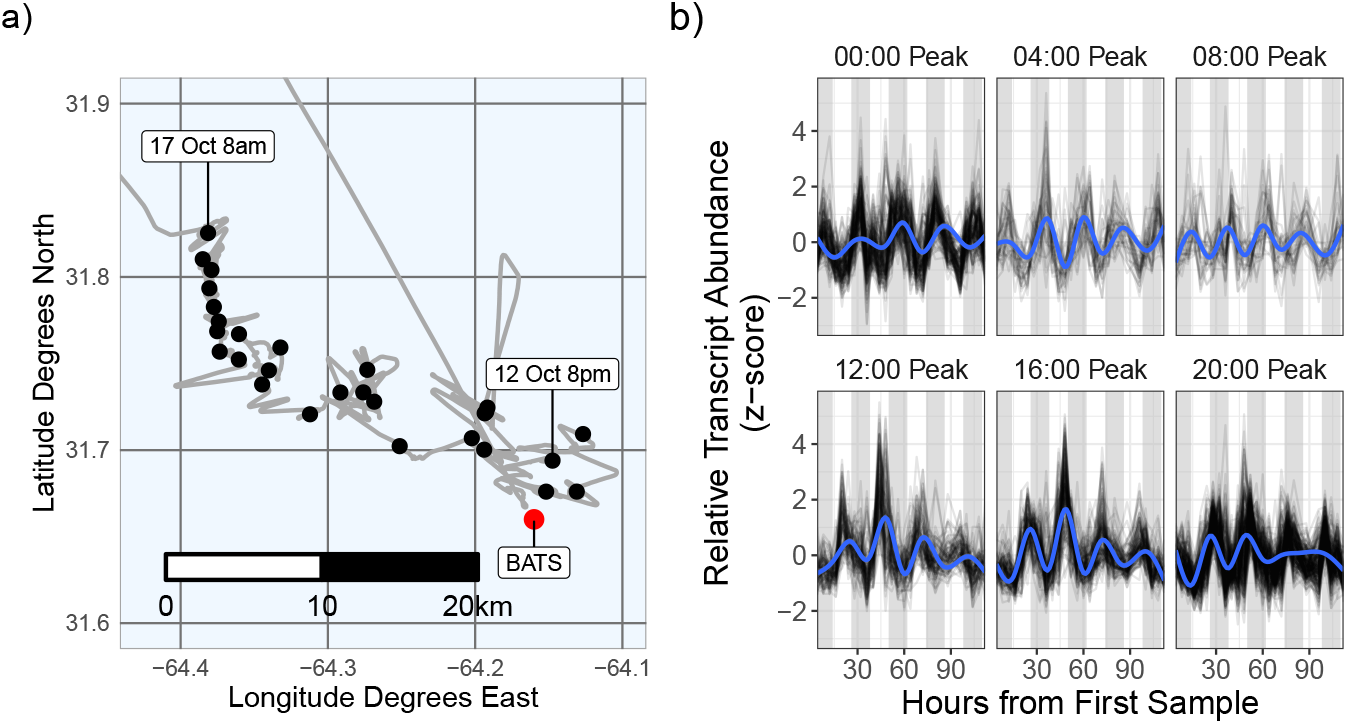
Lagrangian cruise track with BATS station annotated in red and sampling locations annotated in black. Initial and final samples are labelled with local date and time. b) Example gene transcript abundance time series for diel genes annotated as ABC-type transporters. Individual black lines represent the transcript abundance of one gene, the blue line represents a generalized additive model smoothing among all genes. Panels indicate the classification of genes by peak time as determined by the peak time analysis described in Methods. Shaded vertical bars indicate hours after sunset and before sunrise (18:00-06:00 local time).

We estimated the peak time of day for each gene determined to have diel periodicity. Eukaryotic phytoplankton, cyanobacteria, and heterotrophic bacteria had diel expression patterns of different genes occur throughout all times of day, with eukaryotic phytoplankton expressing the greatest number of diel genes at dusk, heterotrophs during the day, and cyanobacteria divided between dawn and dusk (SI Data S3). Example gene expression time series for diel genes with annotations as ABC-type transporters are shown in Figure 1b.

Next we assessed the extent to which the diel timing was synchronized among genes with similar putative function. Genes were functionally annotated using the Kyoto Encyclopedia of Genes and Genomes (KEGG) orthology assignments [1]. The diel genes spanned 3,647 ortholgues. We further analyzed the 849 KEGG orthologues with diel expression in at least four different taxa (eukaryotic phyla/cyanobacterial genera/heterotrophic bacterial orders). For each of these orthologues, we calculated the mean difference in peak time between all pairs of taxa with diel expression of that orthologue. We used an empirical hypothesis testing framework to assess whether that difference was significantly smaller than would be expected by chance given the data set (see Methods). At a false discovery rate of 10%, we identified 324 out of 849 orthologues with significantly synchronized timing across taxa. Significantly synchronized genes included cell division and replication related functions (*ftsH,dnaK* ), photosynthesis-related genes (*psbD,psbA*), and carbon metabolism genes related to the tricarboxylic acid (TCA) cycle and glycine metabolism (*sdhA,coxA*) (Figure 2, SI Data S4).

**Figure 2.**
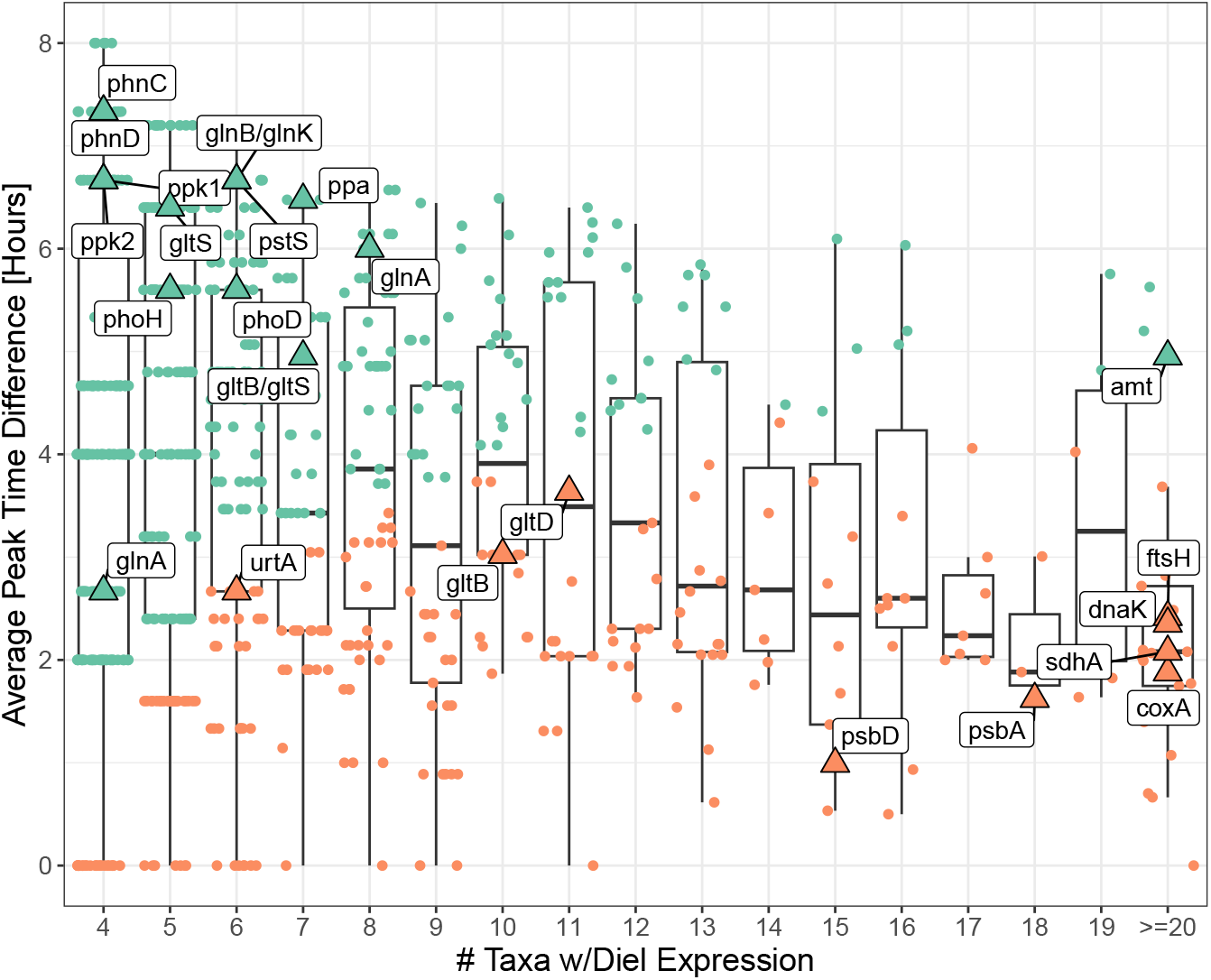
Mean pairwise differences in peak timing for genes with diel expression across at least 4 distinct taxa. X-axis indicates the number of cyanobacterial genera/heterotrophic bacterial orders/eukaryotic phyla with diel expression of a particular KEGG Orthologue as indicated by the periodicity analysis described in Methods. Y-axis indicated the average difference in hours of the peak time of expression for each pair of orthologues (0 means all taxa have peak expression at the same time of day). Key genes in nitrogen assimilation and phosphorus assimilation are indicated. Orange point color indicates a gene with significantly (FDR=10%) synchronized transcription. Synchronization was determined using a Monte Carlo based simulation of the distribution of peak times observed in the full dataset, more details in Methods and detailed description of statistical framework in [28] (also see Supplemental Data 4). Green points indicate a gene without synchronized transcription, labeled ‘asynchronous’.

A pattern also emerged from genes with high degrees of asynchrony across taxa. In particular, genes encoding the uptake and assimilation of nutrients had high asynchrony. We found asynchronous expression of some nitrogen assimilation-related genes, such as components of the GS-GOGAT system and its regulatory proteins as well as the ammonium *amt* uptake receptor (Figure 2). Asynchronized expression of these genes is temporal niche partitioning, which may help to reduce the severity of competition for ammonium in the NPSG [28]. These results may suggest that this form of nitrogen partitioning is common to oligotrophic oceans. However, we found an even higher degree of asynchrony in genes involved in the uptake and incorporation of phosphorus in the Sargasso Sea study site. The phosphonate transporter components *phnC* and *phnD*, inorganic phosphorus transporter *pstS*, phosphate stress regulatory genes *phoH* and *phoD*, inorganic pyrophosphatase *ppa*, and polyphosphate kinases *ppk1* and *ppk2* had among the highest degrees of asynchrony in the data set (Figure 2). A higher degree of asynchrony for phosphorus uptake functions may be further indication of the relevance of diel temporal partitioning as an emergent ecological strategy to overcome limitation of shared scarce resources.

We then examined the taxon-specific transcriptional dynamics of genes involved in the uptake of inorganic and organic phosphorus species to look for specific evidence of uptake partitioning among different functional groups of organisms. We extracted KEGG orthologues specifically annotated as either inorganic phosphate transporters (*pstABC,pstS* - KEGG orthologues K02036,K02037,K02038,K02040) or organic phosphonate transporters (*phnCDE,phnK,phnS* - KEGG orthologues K02041, K02042, K02044, K05781, K11081) and analyzed the relative peak timings of different taxonomic groups. The transporters *pstS, phnC*, and *phnK* had significantly diel transcript abundances in at least 4 different taxa and were analyzed for synchronicity. None of these genes had synchronized expression across taxa. We then collected the expression timeseries of all phosphorus uptake-related transporters to analyze the differences in expression dynamics between cyanobacteria, eukaryotic phytoplankton, and heterotrophic bacteria (see Figure 3). The inorganic phosphorus transporter gene *pstS* had diel transcription in Rhodospiralles and another alphaproteobacterial order peaking at 08:00 (roughly corresponding to sunrise) along with transcripts assigned to the phylum Dinophyceae, while transcripts assigned to diatoms peaked at 12:00 (noon) and both *Prochlorococcus* and *Synechococcus* had peak expression at 20:00 (roughly corresponding to dusk) (Figure 3).

**Figure 3.**
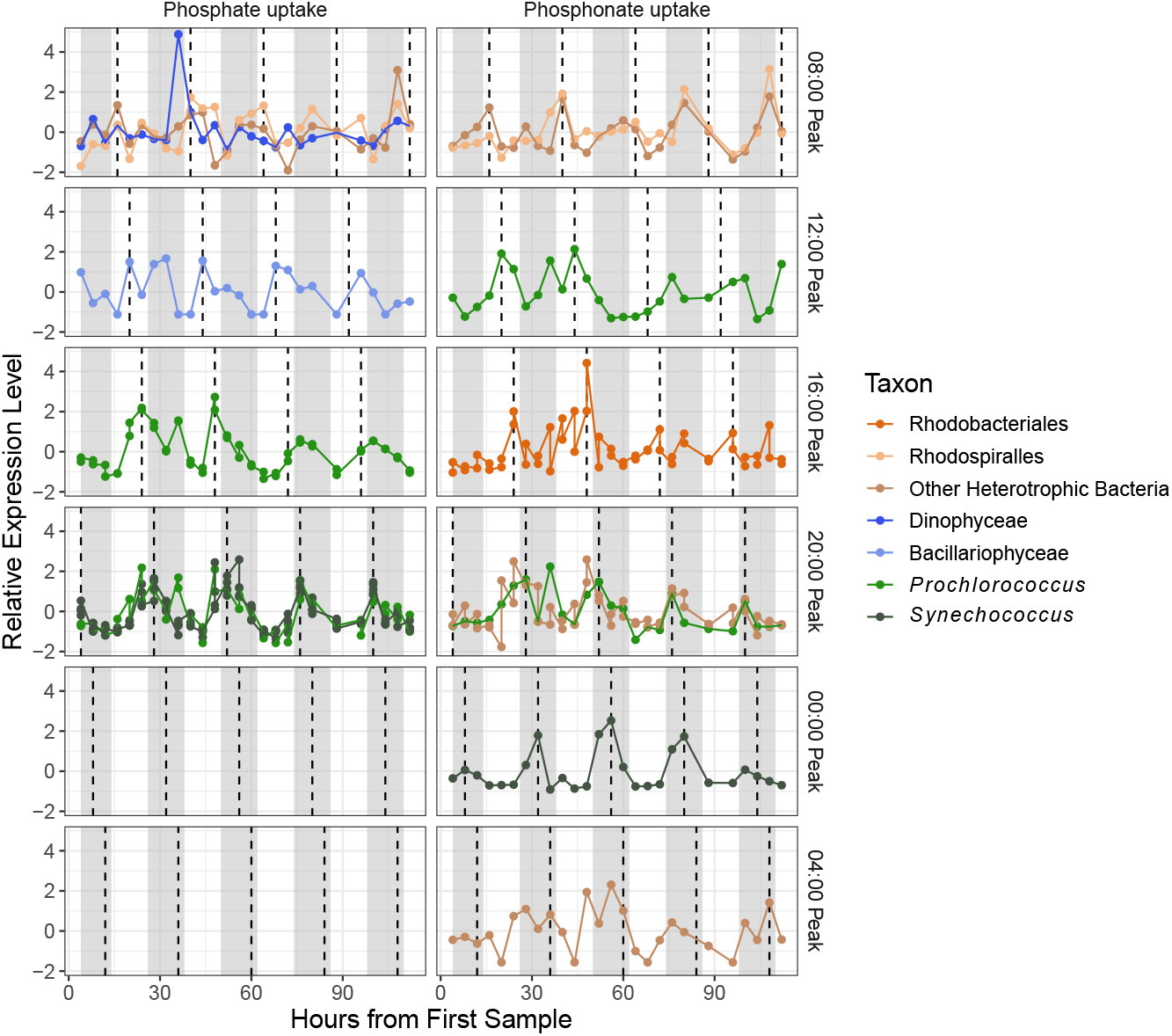
Temporal dynamics of phosphorus acquisition gene transcription. Each line plot shows the gene transcript abundance time series of a specific phosphorus-uptake related gene for a specific eukaryotic phylum or bacterial order (or cyanobacterial genus) identified as diel by the analysis described in Methods and available as Supplemental Data 2. Line color indicates the taxa expressing the transcript. Panels are divided horizontally by time of peak expression as calculated in methods, with 08:00 representing 8am local time, 12:00 noon local time, and so on. Shaded bars indicate hours between sunset and sunrise (18:00-06:00 local). Dashed lines are added to correspond to the peak time of genes in each panel. Panels are divided vertically by the species the gene is involved in transporting. The phosphate uptake category includes phosphate transporter subunits *pstABC,pstS* (KEGG K02036,K02037,K02038,K02040). The phosphonate uptake category includes the phosphonate transporter subunits *phnCDE* and putative phosphatase uptake subunits *phnK,phnS* (KEGG K02041, K02042, K02044, K05781, K11081) - pathway categorization for each gene and calculated peak time are presented in Supplemental Data 3. Panels with no time series plotted indicate that there were no diel phosphate or phosphonate transporters with the labelled peak timing identified in any taxon in the dataset.

Additionally to inorganic phosphate transporters, organic phosphonate transporters are widely transcribed by heterotrophic bacteria, primarily alphaproteobacteria, throughout the day. No eukaryotic organisms had significantly diel transcription of *phn* phosphonate transporters. *Prochlorococcus* and *Synechococcus* had diel expression of the phosphonate transporter subunits *phnC* and *phnD* (*Prochlorococcus* only), but *Prochlorococcus* had peak transcript abundance of these genes during the day and at dusk, concomitant with inorganic phosphorus transporters, while *Synechococcus* expressed *phnC* at 00:00 (midnight), after transcript abundance of *pstS* began to decline. Rhodospiralles transcript abundance of both *pstS* and *phnD* peaked at the same time (08:00). While inorganic phospate transporter transcript abundances were not significantly diel for the alphaproteobacterial order Rhodobacterales, transcripts for the organic phosphonate transporters *phnC* and *phnK* did have diel cycles in abundance, both peaking at 16:00 (Figure 3). Notably, diel cycles of all phosphorus uptake-associated genes within a given taxa tended to peak at the same time of day. The only exception in our dataset was *Synechococcus*, which appeared to exhibit a temporal cascade from primarily inorganic phosphorus transporter transcripts at dusk to phosphante transporter transcripts at midnight. *Synechococcus* polyphosphate kinase genes *ppk1* and *ppk2* also peaked at dusk, concurrently with *pstS* transcripts. Although there is a difference in peak time between *Synechococcus* phosphate and phosphonate transporters, matching timing between *Synechococcus* polyphosphate kinase and phosphate transporters further suggest that *Synechococcus* mainly assimilated phosphate into polyphosphates at dusk rather than assembling polyphosphates from degraded phosphonates. Apart from *Synechococcus*, the timing of diel expression for phosphorus uptake transporters is not influenced by phosphorus being in organic versus inorganic forms. These results suggest that diel partitioning of phosphorus uptake is not entirely due to extrinsic diel cycles in phosphorus speciation, e.g. photolysis of phosphorus-containing dissolved organic matter.

## Discussion

In this study, we tested the hypothesis that microbial ecosystems would temporally partition phosphorous uptake and utilization gene expression in a phosphorus-stressed open ocean oligotrophic ecosystem in the North Atlantic, paralleling previous findings on nitrogen uptake from a primarily nitrogen-limited ecosystem in the North Pacific [28].

Using community metatranscriptome data from a Lagrangian field campaign in the Sargasso Sea, we identified diel gene transcription of phosphorus uptake and assimilation genes across diverse microorganisms. We found phosphorus uptake transporter expression patterns were not synchronized across taxa. Genes associated with organic and inorganic phosphorus uptake and assimilation had higher measured asynchrony in peak timing across taxa than genes related to nitrogen uptake and assimilation. Examining the dynamics of phosphate and phosphonate transporters in more detail, we identified a diel cascade in phosphate transporter expression starting with heterotrophic bacteria in the morning, eukaryotic phytoplankton during the day, and cyanobacteria at dusk. Likewise, we found diel gene transcript abundances for phosphonate transporters with the heterotrophic bacterial order Rhodospiralles expressing phosphonate transporters in the morning (coinciding with phosphate transporters), Rhodobacter during the day, *Prochlorococcus* at dusk (coinciding with its phosphate transporters), and *Synechococcus* overnight (following its phosphate transporters).

Collectively, these results suggest temporal partitioning of uptake for both phosphorus species, and coinciding timing of phosphate and phosphonate uptake on a taxon-by-taxon basis further suggests that substrate-level specialization is not necessarily sufficient to avoid competition for this scarce resource [26], and further suggests diel cycles in transporter gene transcription are not directly tied to changes in the relative pools of different phosphorus species. Complementary evidence for diel temporal niche partitioning for distinct nutrients (N in the North Subtropical Pacific [28] and P in the Sargasso as shown here) may be indicative of complex, community-level metabolic plasticity across ocean basins. Diel temporal niche partitioning to mitigate competition for limiting resources in oligotrophic environments represents an additional mechanism to resolve the paradox of the plankton and the maintenance of complex, marine microbial communities [16].

## Materials and Methods

### Sample Collection and Processing

Fieldwork was conducted from 12 October to 19 October, 2019, in the Sargasso Sea near the location of the BATS station. A drogue was deployed at a depth of 30m (within the surface mixed layer) before the first cast to facilitate a Langrangian sampling strategy. Water column sampling took place every 4 hours for a period of approximately 5 days sampling 5m depth.

### Gene Expression Diel Periodicity and Synchronization Analysis

Please see Supporting Information for more details on timeseries analysis. Briefly, variance-stabilized RNA read counts were subjected to RAIN nonparametric diel analysis as described in [28]. Peak rank time calculations were performed and synchronicity analysis was conducted as described in [28].

### Metatranscriptome Preparation, Sequencing, and Processing

Seawater was filtered through a 0.2 micron pore size Sterivex filtration unit and immediately transferred to a −80^*°*^ C freezer. RNA was extracted using a publicly available phenol-chloroform based protocol [12] and DNA was reduced using the Turbo DNA-free kit (Ambion). Metatranscriptome libraries were prepared by reducing ribosomal RNA using the QIAGEN’s FastSelect kit (Bacteria) and sequenced (2 x 151 nt) using the low-input protocol for total RNA on the Illumina Novaseq S4 platform under the DOE Joint Genome Institute (JGI) Community Sequence Proposal ID 505733. For samples with *<*10 ng of RNA total, the ultra-low input protocol was used. Raw read filtering and trimming was done using BBDuk v38.67 and BBMap v38.84 from the BBtools packages [5]. Trimmed filtered reads were concatenated across samples and assembled using MEGAHIT v1.2.9 [22]. MetaGeneMark v3.38 was used to call open reading frames (ORFs). For the cellular community, trimmed filtered reads were mapped to the combined assembly using BBMap v38.84 [5] using default parameters and tabulated using featureCounts [23]. The ORF protein sequences were annotated using eggNOG-mapper v2.1.4 (with DIAMOND blastp alignment [15]) for functional annotation, and aligned to the PhyloDB database (https://github.com/allenlab/PhyloDB) using the software package EUKulele [18] for taxonomic annotation.

### Diel Transcript Abundance Analysis

Prior to timeseries analysis, all metatranscriptome read mappings were transformed using the DESeq2 variance stabilizing transformation [25]. Methods for periodicity determination and peak time estimation are described in detail in [28]. Briefly, for each time series a linear trend was removed to account for moving averages. Then, the rain [36] method was applied and an FDR control at 10% were set to assess significance. For significantly diel time series, peak time was estimated by calculating the mean rank of each time of day across all samples and selecting the largest.

### Transcript Synchrony Analysis

Methods are described in detail in [28]. Briefly, only KEGG orthologues with diel transcription in at least 4 different taxa were analyzed. For each orthologue, the pairwise difference in estimated peak time was calculated for each pair of taxa with diel expression of that orthologue. The mean of this value across all pairs was used as a test statistic and significance was assessed using a monte carlo simulated null ensemble of all peak times from the dataset. Significance was set at an FDR=10% threshold.

## Data and code availability

Sequence data are publicly available under the DOE JGI Community Sequencing Proposal ID 505733 (https://genome.jgi.doe.gov/portal/Infvirtimeseries/Infvirtimeseries.info.html) Code and processed sequence data used for analyses are available at https://github.com/d-muratore/phosphorus_partitioning.

Supplemental files available for this manuscript are described below:

### Supplemental Data 1

Ship navigation data with sampling locations and sample environmental data for all metatranscriptome samples.

### Supplemental Data 2

Results of RAIN diel periodicity analysis. Columns include KEGG orthologue tested, taxonomic assignment of KEGG orthologue, RAIN test p-value, and a reject/fail to reject decision on the null hypothesis that the transcript does not have a 24-hour diel oscillation.

### Supplemental Data 3

Calculated peak time, assigned taxnomic affiliation, and KEGG orthologue pathway category for all transcripts determined to be significantly diel, as designated in Supplemental Data 2.

### Supplemental Data 4

Results of KEGG orthologue synchronicity analysis. Columns show the KEGG Orthologue number of the gene tested, the number of taxa in the dataset with diel expression of this KO, the pathway assignment of the KO, the average difference in peak rank time between all organisms with diel expression of the KO, the associated empirical p-value (10,000 Monte Carlo simulations) and the reject/fail to reject decision of the null hypothesis that the transcript is not synchronized.

## Acknowledgments

The authors would like to thank the captain and crew of the RV Atlantic Explorer for conducting cruise AE1926. This work was funded by NSF grant OCE-1829641 to S.W.W., NSF grant OCE-1829636 to J.S.W., Simons Foundation grant 721231 to J.S.W., and the Blaise Pascal Institute Chair of Excellence award at the Institut de Biologie of the École Normale Supérieure to J.S.W. The work (proposal: 10.46936/10.25585/60001286) conducted by the U.S. Department of Energy Joint Genome Institute (https://ror.org/04xm1d337), a DOE Office of Science User Facility, is supported by the Office of Science of the U.S. Department of Energy under contract no. DE-AC02-05CH11231. We thank Christine Sun and Matthew Sullivan for their assistance as co-investigators on contract no. DE-AC02-05CH11231. Work at LLNL was performed under the auspices of US Department of Energy Office of Science contract DE-AC52-07NA27344. Institution Paper Number LLNL-JRNL-860896-DRAFT.

